# VDJdive and ECLIPSE enhance single-cell TCR sequencing analysis through the probabilistic resolution of ambiguous clonotypes

**DOI:** 10.64898/2026.02.18.706444

**Authors:** Ethan C Burns, Mercedeh Movassagh, Jill F Lundell, Mingzhi Ye, Zhaochen Ye, Giacomo Oliveira, Rishabh Rout, Miya B Hugaboom, Kelly Street, David A Braun

**Affiliations:** Department of Immunobiology, Yale School of Medicine, New Haven, CT, USA; Center of Molecular and Cellular Oncology, Yale Cancer Center, Yale School of Medicine, New Haven, CT, USA; Department of Neurosurgery, Yale School of Medicine, New Haven, CT, USA; Sonic Healthcare USA, Dallas, TX, USA; Harvard T.H. Chan School of Public Health, Boston, MA, USA; MIT Lincoln Laboratory, Lexington, MA, USA; Division of Biostatistics, Keck School of Medicine, University of Southern California, Los Angeles, CA, USA; Program in Computational Biology and Biomedical Informatics, Yale University, New Haven, CT, USA; Department of Medical Oncology, Dana-Farber Cancer Institute, Boston, MA, USA; Harvard Medical School, Boston, MA, USA; Section of Medical Oncology, Department of Medicine, Yale School of Medicine, New Haven, CT, USA; Department of Pathology, Yale School of Medicine, New Haven, CT, USA; Department of Urology, Yale School of Medicine, New Haven, CT, USA

## Abstract

Single-cell T cell receptor sequencing (scTCR-seq) has transformed our ability to track individual T cell clones and has been instrumental in advancing our understanding of human T cell differentiation. However, current computational pipelines for analysis, which require precise matching of CDR3 sequences from exactly 2 heterodimeric TCR chains to define the clonotype for each cell, are inherently limited because of the substantial proportion of cells possessing “ambiguous” clonotypes driven by missing (undetected from the technical issue of chain “dropout”) or extra chains (present from either true biological expression or due to technical artifacts such as cellular doublets and ambient TCR contamination). As a result, clone sizes are artificially reduced, impeding the tracking of clones across conditions and differentiation states. Here we introduce VDJdive and ECLIPSE (Enhanced CLonotypic Inference via Prediction of Single-cell Expression), two computational methods that, together, resolve this clonal ambiguity by utilizing the expectation-maximization algorithm for the clonal prediction of ambiguous cells. These methods consider chain pairings across the sample, allowing for high-fidelity prediction of chains lost due to dropout and the discernment of biological expression of extra chains from technical artifacts. Consequently, clone sizes are augmented and cells without clonotype assignments are minimized. Our approach facilitates enhanced clonal tracking through these elevated clone sizes and is easily implementable, compatible with standard single-cell transcriptomic workflows, and broadly applicable across biological contexts and T cell subsets.

## Introduction

T cells are an essential arm of the adaptive immune system that is tasked with performing proteomic surveillance throughout the body. Central to this ability are heterodimeric T cell receptors (TCRs) which define the antigen specificity of each clone and are created through the process of V(D)J recombination. Because this process can generate up to 10^18^ unique αβ TCRs, TCRs are also unique to each clone, allowing them to serve as a molecular barcode for the identification and tracking of cells of each clone. Furthermore, TCRs influence the propensity of each T cell differentiation trajectory (Simoni et al., 2018; Oliveira et al., 2021; Abdullah et al., 2025). Clonotypes and their respective TCR must therefore be considered in studies of T cell differentiation and function for a comprehensive understanding of these processes. This approach has proven fruitful in cancer immunology, where clonal tracking has revealed that the clones that differentiate into terminally differentiated cytotoxic effector cells are distinct from those linked with a dysfunction/exhausted phenotype (Li et al., 2019, Zhang et al., 2018). Similarly, clonal tracking has been crucial in establishing that both murine and human tumor-draining lymph nodes maintain a reservoir of stem-like CD8+ T cells that become progressively dysfunctional within the tumor microenvironment (Connolly et al., 2021, Prokhnevska et al., 2023).

Traditionally, the tracking of T cell clones was a challenging process, but the advent of single-cell RNA sequencing with paired TCR sequencing (scRNA/TCR-seq) has dramatically increased cellular throughput and speed. Because this assay recovers both the TCRα and the TCRβ, the conventional method of defining TCR clones has been by the pairing of the most diverse sequence (either amino acid or RNA) of each chain: the CDR3 region. However, “dropout” of transcripts is common in scRNA-seq, whereby only a fraction of RNA transcripts present in each cell end up being detected by the assay (Macosko et al., 2015; Qiu, 2020). This can result in only 1 or no TCR chains being detected in a sizable proportion of cells (Borcherding et al., 2020; Sturm et al., 2020). The cells of this population are clonally “ambiguous” as a result, causing many analysis pipelines to filter them out and substantially reducing the number of cells that can be analyzed (Le Coz et al., 2023; Zheng et al., 2021; Perez et al., 2022). An alternative approach has been to define these ambiguous cells as being in clones marked by the presence of 1 specific chain alongside the absence of all others (Minowa et al., 2024). While this maximizes the proportion of cells able to be retained, it artificially restricts clone sizes and inflates TCR diversity.

An additional complexity of TCR analysis is the phenomenon that in many cells, 3 unique TCR chains are recovered (Zhu et al., 2023). These cells are often excluded from analysis or one chain is artificially removed (typically based on expression level) to force a 1:1 pairing (Sturm et al., 2020; Muhowski and Rogers, 2023). The common rationale for their removal is the possibility that these additional detected TCR chains represent technical artifacts that are inherent to scRNA/TCR-seq, namely ambient TCR contamination and cellular doublets (Young and Behjati, 2020; Zheng et al., 2017). Additionally, their presence would seem to defy Burnet’s clonal selection theory as the addition of a third chain could convey more than one specificity to each clone (Burnet, 1976). Despite this, it has been described for several decades that T cells can biologically express an extra TCR chain, either at the RNA or protein level (Malissen et al., 1988; Matis et al., 1988; Triebel et al., 1988; Padovan et al., 1993; Ji et al., 2010). The presence of an extra chain is more common for TCRα than with TCRβ, with some estimates predicting ∼10% of cells have an extra alpha TCRα chains and ∼1-7% have an extra TCRβ chain (Muhowski and Rogers, 2023). Other studies have supported even higher frequencies, including even up to one-third of all T cells (Yang et al., 2020; Padovan et al., 1993; Carter et al., 2019). Therefore, the automatic removal of cells with extra TCR chains eliminates biologically relevant cells and impedes the ability to effectively track T cell clones and assess TCR diversity.

To address these challenges, we developed VDJdive and ECLIPSE. First, VDJdive implements the expectation-maximization (EM) algorithm to resolve clonal ambiguity in cells with extra or missing TCRs by considering the chain pairings of other cells present in the sample. Next, ECLIPSE (Enhanced CLonotypic Inference via Prediction of Single-cell Expression) utilizes VDJdive in a user-friendly, Seurat-compatible R package that explicitly accounts for the biology where T cells can harbor extra TCR chains (Hao et al., 2023). ECLIPSE generates Seurat objects in a format that is compatible with downstream analysis with the comprehensive scRepertoire package, allowing for streamlined visualization and diversity analysis (Borcherding et al., 2020; Yang et al., 2025). By addressing the technical and biological complexities of scTCR-seq, VDJdive/ECLIPSE augment T cell clone sizes and reduce missing data without sacrificing the phenotypic identity of each T cell clone. Our approach facilitates analysis that would not be possible with smaller clone sizes and is broadly applicable across T cell subsets and sample contexts.

## Results

### Clonal ambiguity is prevalent in scTCR-seq data

Given that a standard approach to scTCR-seq analysis is to include only cells with exactly 1 TCRα and 1 TCRβ, we first investigated what proportion of cells meet these strict criteria. To address this question, we utilized an existing scRNA/scTCR-seq dataset of paired tumor and adjacent normal tissue samples from 13 patients with clear cell renal cell carcinoma (ccRCC) (Braun et al., 2021). This dataset contains 65,258 CD8+ T cells across 13 clusters, falling into 5 broad phenotypes: cytotoxic, proliferating, resident memory (T_RM_), exhausted, and terminally exhausted (**Fig. 1a**).

**Figure 1:**
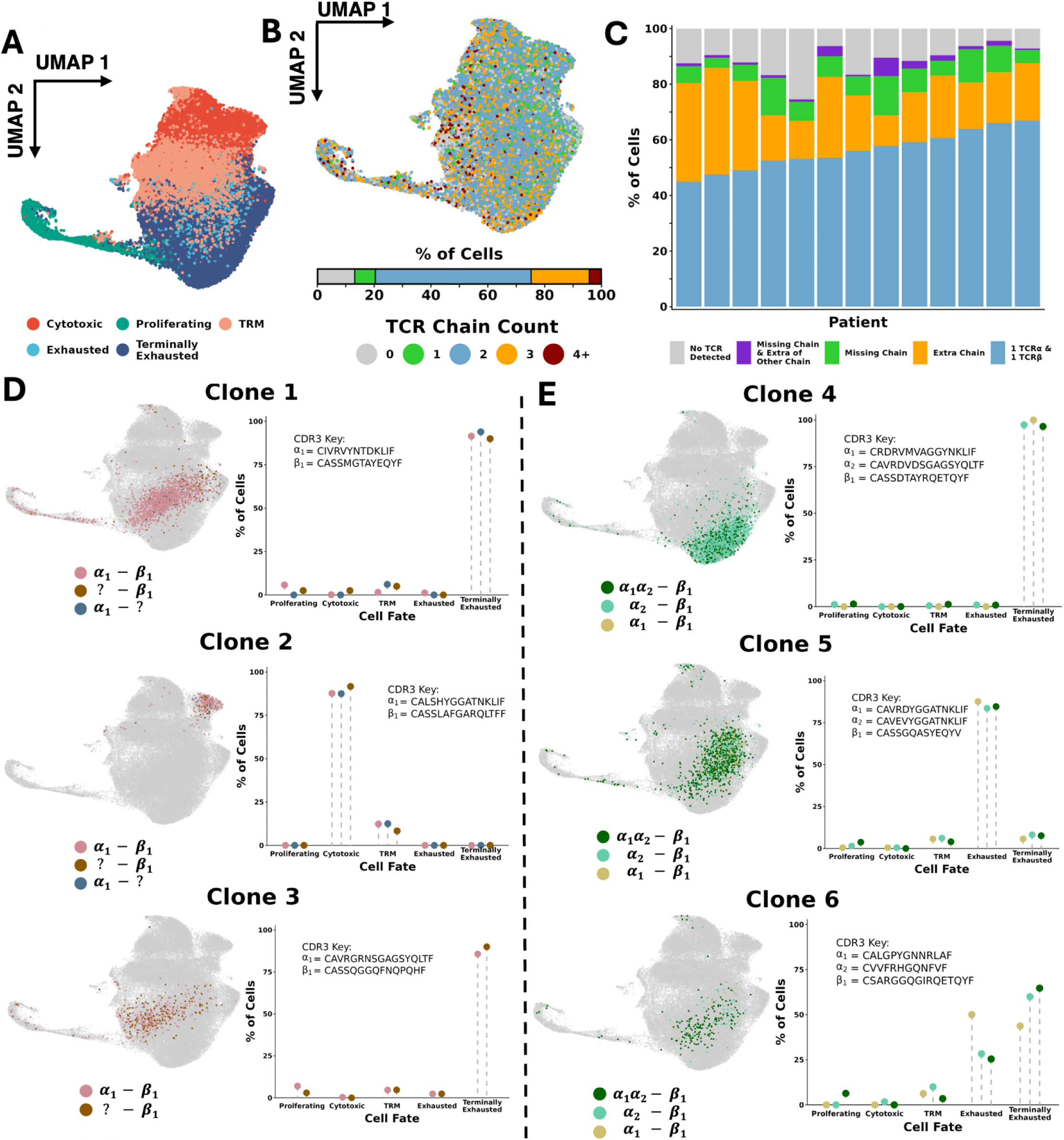
Cells with ambiguous clonotypes are prevalent m scTCR-seq and clonally related to cells with resolved clonotypes. A. UMAP of CD8+ T Cells in testing (renal cell carcinoma) dataset. 5 distinct cell fates are shown, originating from 13 original clusters. B. Number of total TCRa and TCRp chains detected per cell. C. Chain combinations of TCRa and 1 TCRp detected per cell, then split by patient. D. 3 putative clones are highlighted, each possessing some cells with only TCRp, others with that TCRp and a TCRa, and others with just the paired TCRa. On the left, the cells are shown projected onto the UMAP of all CD8+ T cells and colored depending on the chains that were detected. On the right, the percentage of cells with each chain configuration of each cell fate is shown. E. Identical to E, except the potential clones all may possess an extra TCRa.

Mapping of the number of unique TCR chains (TCRα + TCRβ) detected per cell revealed that, strikingly, only 54.8% of cells expressed exactly 2 chains (**Fig. 1b**). 20.4% of cells expressed 1 or no TCR chains, while another 20.4% of cells expressed 3 TCR chains. When split by chain type, the median percentage of cells per patient with precisely 1 TCRα and 1 TCRβ was 56.0% (range: 44.9 - 66.9%) (**Fig. 1c**). In contrast, the median percentage of cells that expressed a set of at least 1 TCR chain that didn’t fit this pattern was 30.0% (range: 21.4 – 42.9%). This indicates that clonally ambiguous cells lacking a 1:1 α-β TCR pairing represent a substantial proportion of cells in scTCR-seq. Therefore, removing these cells with fewer than or more than 2 chains (30.0% of cells) would exclude a large percentage of cells from analysis. In contrast, their inclusion could greatly enhance downstream analysis as far more cells defined at the RNA level would be assigned a matching clonotype.

### Clonally ambiguous cells are phenotypically similar to clonally resolved cells

Due to the appreciable frequencies of clonally ambiguous cells (possessing a missing or extra TCR), we sought to elucidate the true clonotype of these cells to prevent their exclusion from analysis. One strategy to address this is to predict the true clonotype by leveraging the 1) detected TCR chains of each ambiguous cell, and 2) the frequency of clonally resolved cells with shared chains and a strict 1:1 α-β TCR pairing. We hypothesized that if clonally ambiguous cells are truly clonally related to clonally resolved cells, then these populations should share a similar phenotype. This is because cells derived from the same T cell clone originate from a shared progenitor and are influenced by shared antigen exposure, tissue microenvironments, and differentiation patterns, ultimately leading to similar functional states and phenotypes (Simoni et al., 2018; Oliveira et al., 2021; Abdullah et al., 2025). To investigate this hypothesis, we examined three expanded sets of cells that all expressed solely a TCRβ chain (clonally ambiguous) and compared their phenotypes to clonally resolved cells (1:1 α-β TCR pairing) that share the same TCRβ CDR3 amino acid sequence. In all three instances, the clonally ambiguous cells shared the same phenotype(s) as the clonally resolved cells, supporting the hypothesis that they are part of the same clone (**Fig. 1d**). Further, for 2 of the 3 potential clones, there was an additional set of clonally ambiguous cells that displayed only the TCRα chain of the clonally resolved cells. Consistent with this hypothesis, the “α-only” clonally ambiguous cells shared the same phenotype as both the clonally resolved cells and the “β-only” clonally ambiguous cells. Therefore, across these 3 putative clones, the phenotype was consistent regardless of which TCRα or TCRβ was detected, while remaining highly specific to the individual clone being examined.

We next evaluated whether clonally ambiguous cells with an extra TCR chain are also phenotypically similar to clonally resolved cells. If clones truly expressed 3 TCR chains *in vivo* (i.e. biological expression of 3 chains and not a technical phenomenon), then we would expect numerous cells to express the exact same TCR chain pairings. In contrast, random ambient TCR contamination or doublet formation would be highly unlikely to lead to the exact same combination of 3 TCR chains being present together in multiple cells. Analysis of chain pairings revealed numerous instances of multiple cells with identical sets of 3 TCR chains (typically two α chains and one β chain). Importantly, these clonally ambiguous cells with 3 TCR chains (α_1_α_2_-β_1_) shared the same phenotype(s) as the clonally resolved cells with a strict 1:1 pairing (either α_1_-β_1_ or α_2_-β_1_), reinforcing the notion that they all belong to the same clone (**Fig. 1e**). Together, these results support the hypothesis that clonally ambiguous cells belong to the same clonotype as clonally resolved cells within the same sample and suggest that leveraging clonal information across a sample could further resolve clonal ambiguity.

### Development of VDJdive and ECLIPSE to address clonal ambiguity

Having conceptually formed a strategy for the clonal prediction of clonally ambiguous cells, we sought to generate a method that applies the intuition into a rigorous statistical model. To do this, we created VDJdive which implements the EM algorithm to probabilistically assign clonotypes to clonally ambiguous cells (see Methods) (Dempster, Laird, and Rubin, 1977; Ceppellini et al., 1955). Importantly, it operates under the assumption that cells should express one of each chain type, with each clonotype being defined by pairing the CDR3 amino acid sequences of both chains. The clonal assignment of ambiguous cells is then done via the proportional pairing of each cell to clones in a manner that reflects the probability of each potential clone being correct. While VDJdive substantially ameliorates clonal ambiguity for cells with just 1 TCR chain, its assumption that cells should always have exactly 2 TCR chains prohibits clones that biologically express 3 chains from being preserved.

To address this complexity of scTCR-seq, we further developed ECLIPSE (Enhanced CLonotypic Inference via Prediction of Single-cell Expression) which maintains the statistical foundation of VDJdive while explicitly considering the possibility that 3 TCR chains are expressed in some clones (**Fig. 2a**). Under ECLIPSE, clones are permitted to have 3 TCR chains only if 3 or more cells are observed with the identical set of chains, as it is highly unlikely that technical artifacts (such as doublet formation) would produce the same exact TCR chain combination across multiple cells (see Methods for details and additional filters). Clones that are annotated as having 3 chains are also joined with cells that contain 2 of the 3 chains (with 1 TCRα and 1 TCRβ), which mitigates the impact of technical chain dropout. After clones with 3 TCRs are considered, all remaining ambiguous cells are mapped by VDJdive. Rather than proportional assignment, cells are solely assigned to clones if the probability is high enough (generally p ≥ 0.8, see Methods). This results in the joining of cells seen in **Fig. 1d and in Fig. 1e**, as well as other sets of cells if considered statistically probable. For cells that do not meet these probability thresholds, their final clonotype is only defined by the observed chains, and no clonotypic prediction is performed. ECLIPSE therefore probabilistically resolves clonal ambiguity while maintaining clones that express 3 TCR chains. Moreover, our method was also designed to operate with Seurat objects and purposefully formats them for downstream analysis with the comprehensive scRepertoire R package, enabling integration with current standard tools and workflows (Hao et al., 2023; Borcherding et al., 2020).

**Figure 2:**
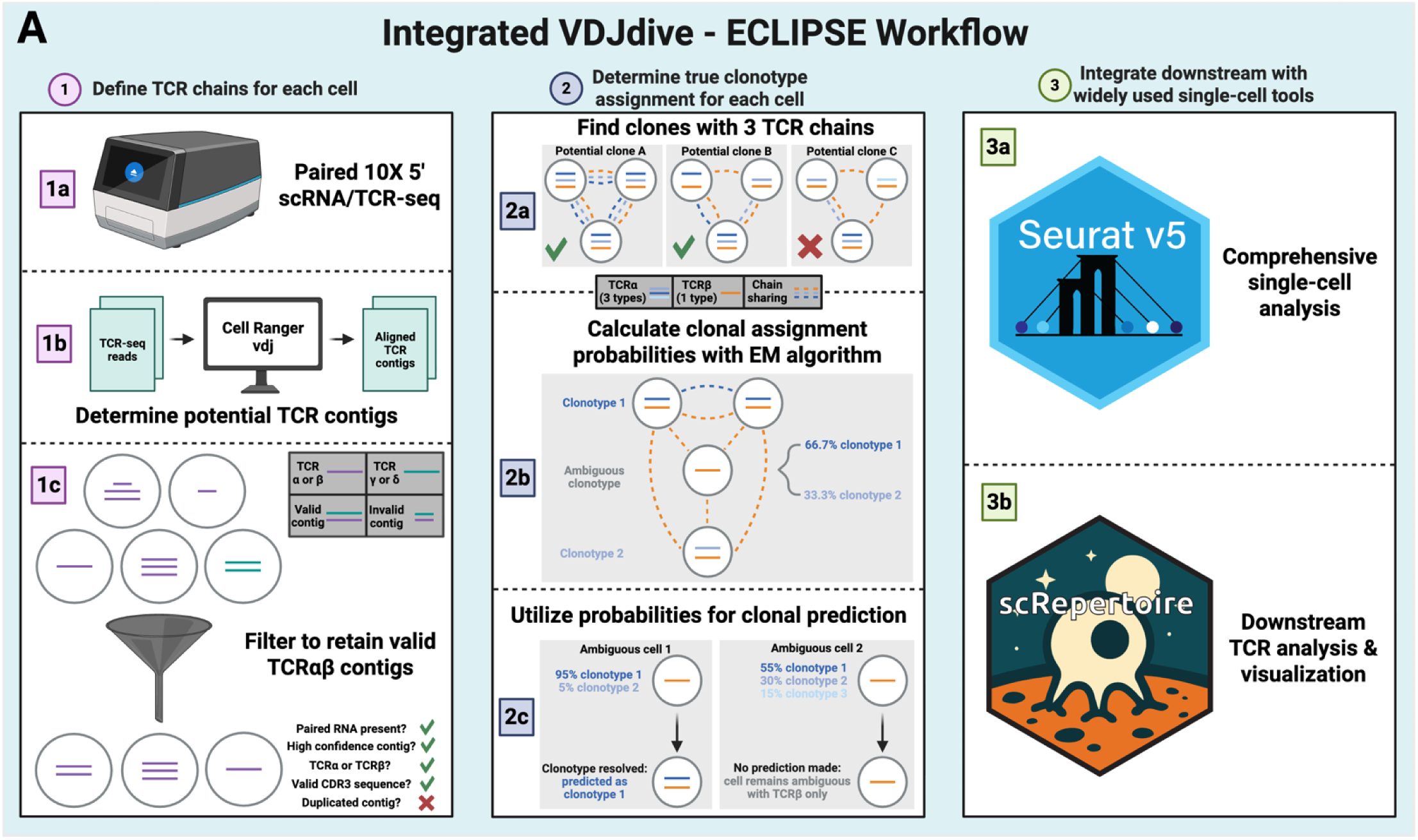
The integrated workflow of VDJdive and ECLIPSE for defining clonotypes with scTCR-seq. A. The integrated workflow is split into 3 parts: defining TCR chains, determining clonotypes, and integrating the results with other tools. Briefly, TCR chain definition is done by performing paired scRNA/TCR-seq, with the reads being aligned with Cell Ranger vdj before chain filtering is performed. These chains are then compared among all cells of each sample to predict which clones have 3 TCR chains. The EM algorithm is then used to predict the probability of each clonotype for each clonally ambiguous cell. The true clonotype of ambiguous cells is then predicted only of the probability of any one clone is high enough. After the clonotype of each cell is defined, the results are integrated into Seurat for comprehensive single-cell analysis. Additionally, the Seurat object is formatted for downstream visualization and clonal analysis with scRepertoire.

### VDJdive and ECLIPSE resolve clonal ambiguity and improve clonotype detection

With VDJdive/ECLIPSE in hand, we first asked whether they could substantially diminish clonal ambiguity and preserve clones with 3 TCR chains. After implementing ECLIPSE, the vast majority of T cells (80.5%) were clonally resolved (**Fig. 3a**). As intended, ECLIPSE also preserved clones with 3 TCR chains (**Fig. 3b**), which accounted for a median of 20.0% of cells (range: 7.3 – 57.9%) per patient. Across patients, the predicted frequency of cells with extra TCRα was higher than that of TCRβ (**Fig. 3c),** with their frequency being positively correlated (**Fig. 3d**).

**Figure 3:**
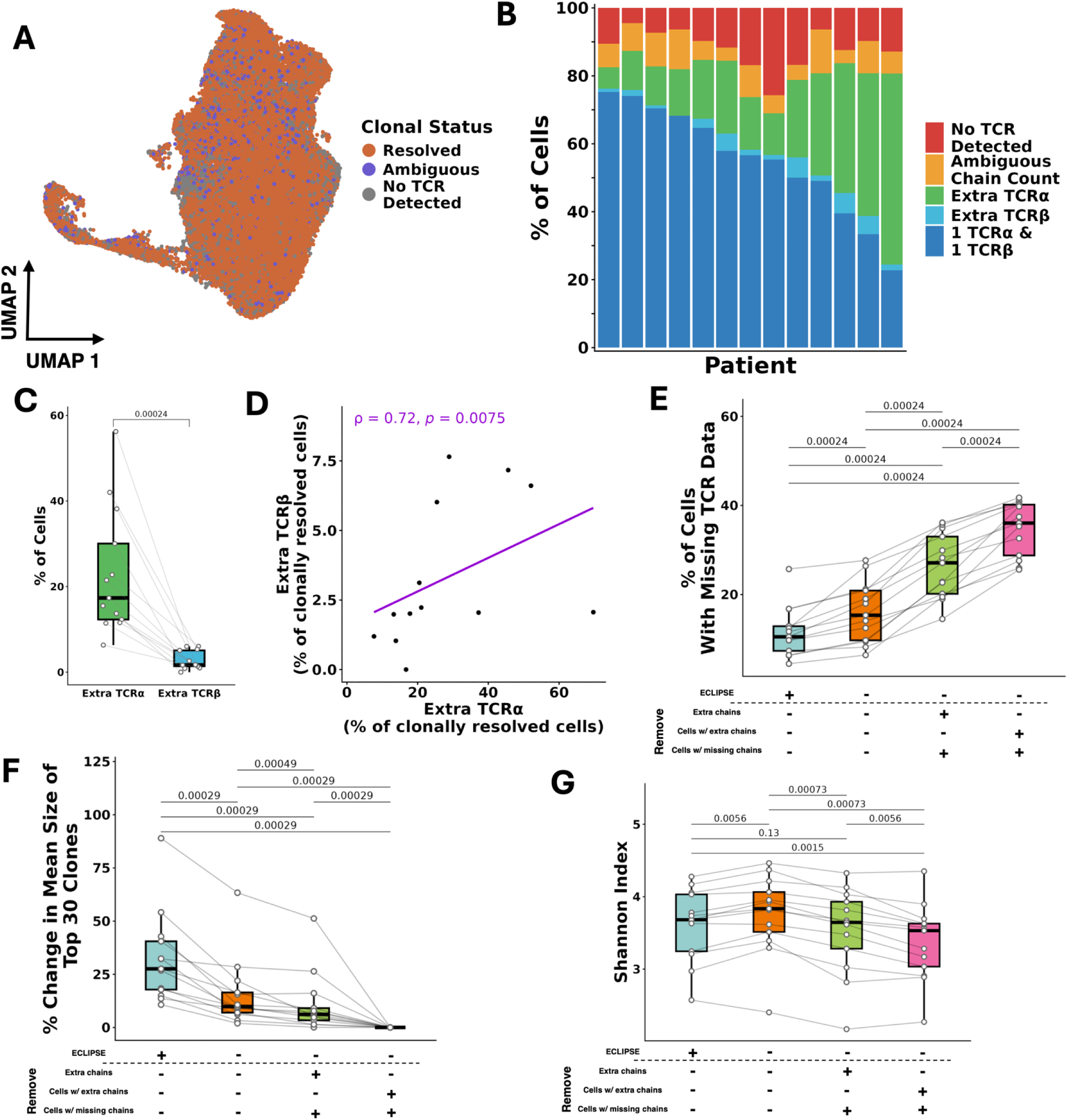
VDJdive/ECLIPSE resolve clonal ambiguity to enhance clonal tracking over current benchmarks. A UMAP showing the clonal annotation of each cell, either clonally resolved, clonally ambiguous, or no TCR detected at all. B. Predicted chains by VDJdive/ECLIPSE per cell and then split by patient. C. The predicted percentages of cells per patient with extra TCRa vs. TCRl3. D. Percentage of cells per patient with extra TCRa chains plotted against that of TCRl3. A spearman correlation was performed. E. The percentage of cells with no TCR annotation per patient plotted by TCR analysis method. F. The percentage change in size of the 30 largest clones per patient plotted by TCR analysis method. G. The Shannon index of TCR repertoire diversity per patient plotted by TCR analysis method. Wilcoxon-signed rank test was used for A, E, F, G, with a false-discovery rate (FDR) correction being used for multiple-testing correction. For D, a Spearman correlation was calculated. p-values are shown above all applicable plots.

We next benchmarked VDJdive and ECLIPSE against existing, standard methods to further quantify their performance. For consistency, the three methods chosen for comparison all defined clones via the CDR3 amino acid sequence of both chains, and they have been adopted in custom pipelines as well as existing packages such as scRepertoire (Borcherding et al., 2020; Yang et al., 2025). The first comparator method considers all cells and chains as is; the second retains only the highest expressed TCRα and TCRβ per cell and removes cells with only a single detected TCR chain; the third removes cells with either missing chains or multiple TCRα or TCRβ, leaving only cells with exactly 1 TCRα and 1 TCRβ. After benchmarking against these three methods, we observed that ECLIPSE significantly reduced the percentage of cells without an annotated clonotype, reducing the median unannotated clonotype frequency to just 10.5% (compared to 15.3, 27.1, and 36.0% with other methods) (**Fig. 3e**). ECLIPSE also resulted in a significantly increased clone size for the 30 largest clones when compared to all other methods (**Fig. 3f**). The magnitude of increase across patients over other methods reached as high as a median of 27.6% across all patients (and up to 89.0% for one patient). We observed similar findings for the next 30 largest clones (ranked from 31-60, collective median clone size of 8 cells) (**Extended Data Fig. 1a**), indicating that this effect is not exclusive to hyperexpanded clones. The increase in clone size was accompanied by a change in TCR repertoire diversity as defined by the Shannon index, specifically a reduction in diversity compared to methods that did not remove any cells/chains and an increase compared to those that did (**Fig. 3g**). Notably, the Shannon index is widely used to quantify T cell repertoire diversity across disease contexts, suggesting that differences in clonotype resolution alone can lead to different conclusions about repertoire diversity, even when applied to the same dataset (Chiffelle et al., 2020). Such differences in repertoire diversity are particularly meaningful, as differences between patients have been associated with overall survival in cancer (Hopkins et al., 2018; Keane et al., 2017). Beyond the Shannon index, similar trends were observed with the Inverse Simpson, Normalized Entropy, and Gini-Simpson indices, although they were not always statistically significant (**Extended Data Fig. 1b-d**). Ultimately, these findings suggest that VDJdive and ECLIPSE resolve clonal ambiguity, resulting in a notable improvement in the detection of moderate and large clonotypes compared to existing methods.

### VDJdive/ECLIPSE accurately recover biologically valid TCR clonotypes

Building on the improved T cell clonal reconstruction achieved by VDJdive/ECLIPSE, we next investigated whether the increased clones were biologically valid using a data simulation approach. We computationally removed or added 500 TCR chains to cells with a ground-truth clonotype (i.e. 1 TCRα and 1 TCRβ), enabling direct evaluation of how accurately VDJdive (and subsequently ECLIPSE) recovered the true clonotype (**Fig. 4a-b**). This analysis demonstrated that in both scenarios, VDJdive was able to predict what was added/missing over 80% of the time (81.0% for chain removal and 86.2% for chain addition). In most of the remaining cells, VDJdive did not predict any clone with high confidence, resulting in ECLIPSE instead defining the clone by only the detected chains. Importantly, VDJdive falsely predicted a clonotype with high confidence that did not match the ground-truth clonotype in only 0.4 - 0.6% of cases (2-3 cells of 500). Cells that were not modified during the simulation were predicted identically with or without the modification of the 500 other cells in 99.94 - 99.97% of cells (55,421 - 55,443 cells of 55,457). Together, these two simulations indicate that the resolution of clonal ambiguity by VDJdive/ECLIPSE is done with high accuracy.

**Figure 4:**
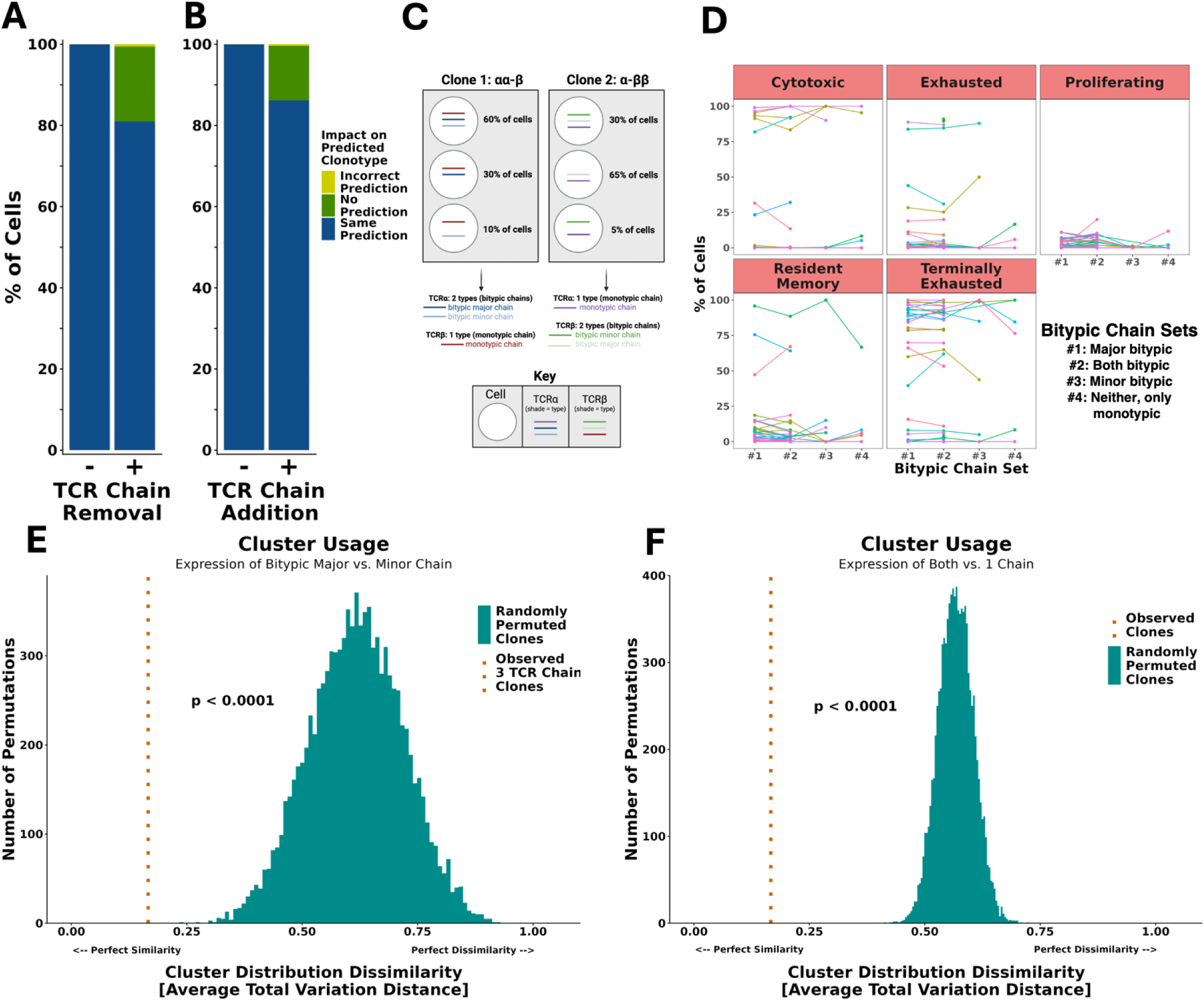
VDJdive/ECLIPSE is able to predict previously ambiguous clonotypes with high accuracy. A. Chain removal simulation, where 500 random contigs were removed from cells with a ground-truth clonotype (i.e. 1 TCR a and 1 p). The final clonal annotation was then compared for these cells with or without the removal of the contig. On the left, unchanged cells were also assessed as a negative control. B. Identical to A, but with the random addition of 500 contigs to 500 random cells. C. Schematic illustrating how bitypic major, bitypic minor, and monotypic chains are identified for each clone with 3 TCR chains. Briefly, the chain with two types (i.e. 2TCRa or 2 TCRP) is called as the bitypic chain, while the chain with one type is defined as monotypic. Among the 2 bitypic chains, the type observed in more cells is labeled as the bitypic major chain, while the other bitypic chain that is observed in fewer cells is labeled as the bitypic minor chain. D. For large clones, the percentage of cells per bitypic chain set of each cell fate. Each line/color indicates a large clone, with the x-axis showing different detected chain sets within the clone. All clones shown here were predicted to contain 3 TCR chains. E. Permutation test of 10,000x randomly permuted sets of groups of randomly paired clones. The mean cluster distribution dissimilarity among the set is plotted (as measured by mean total variation distance). For the observed, all cells were of clones predicted to have 3 TCR chains, with the comparison being whether the bitypic major vs. minor chain was expressed. F. Identical to E, but the comparison was cells with 1+ of each chain type, or not.

As an orthogonal approach to validate the biological faithfulness of recovered clones, we hypothesized that cells assigned to a clonotype by ECLIPSE should closely mirror the phenotypes of other cells within that same clone (i.e. the phenotype of cells within a clonotype are similar). Beyond investigating clones with only 2 TCR chains (**Fig. 4a-b**), here we analyzed cells within clonotypes that VDJdive/ECLIPSE predicted would have 3 TCR chains (either 2 TCRα and 1 TCRβ, or less commonly 1 TCRα and 2 TCRβ). We termed the TCR chain that had two distinct sequences the “bitypic” chain, while the chain with only one sequence was defined as the “monotypic” chain (e.g., for a clonotype that had 2 TCRα and 1 TCRβ, the TCRα would be the bitypic chain and TCRβ would be the monotypic chain) (**Fig. 4c**). In this example, if an individual cell expressed both TCRα chains (regardless of whether TCRβ was also detected), it would be labeled as expressing “both bitypic” chains. If a cell only expressed one of the two TCRα chains, and it was the more commonly expressed of the two TCRα chains, then it would be labeled as expressing the “major bitypic chain”. Conversely, if a cell expressed only one of the two TCRα chains but it was the less commonly expressed TCRα chain, then it would be labeled as expressing the “minor bitypic chain.” Cells that did not express either bitypic chain and thus only contained the monotype chain were marked as monotypic only. Using this nomenclature, we compared the phenotype of these different cell types within an individual clonotype (**Fig. 4d**). Qualitatively, we observed that essentially all cells within an individual clonotype had a similar phenotype, irrespective of which TCR chain the individual cell expressed. To quantify the likelihood that VDJive/ECLIPSE was joining biologically distinct cells together into a clonotype making this a feature of random chance, we performed permutation testing. Following 10,000 permutations, no permuted set of groups of paired clones resulted in cells as similar to their pairing as cells within VDJdive/ECLIPSE-defined clones (which differed only in which bitypic chain was detected; **Fig. 4e**). Likewise, no set of 10,000 permuted groups of paired clones was as similar as cells within ECLIPSE clones compared based on whether they express both 1+ TCRα and 1+ TCRβ chain or not (**Fig. 4f**). This indicates that clones defined by VDJdive/ECLIPSE are remarkably phenotypically consistent, providing evidence that they are indeed true biological clones. Moreover, these results suggest that while VDJdive/ECLIPSE greatly enhance clone sizes, the accuracy of clonal annotation remains high.

### ECLIPSE-defined clonotypes with 3 TCR chains are not doublets

We next investigated whether technical “dropout” of TCR chains was likely responsible for the different number of TCR chains detected between cells in a clonotype. We would hypothesize that cells containing a smaller number of TCR chains due to technical dropout would likely also have less total RNA detected. Within clonotypes with 3 total TCR chains, we observed that cells that expressed both bitypic chains had increased amounts of RNA (RNA counts) and unique genes (RNA features) detected as compared to cells that expressed only 1 of the bitypic chains (major or minor) or neither (and only a monotypic chain) (**Extended Data Fig. 2a-b**). However, this could be caused by doublets (which have been described to increase RNA levels), so we next investigated whether cells with 3 TCR chains could be doublets. Neither cells with 3 TCR chains in clonotypes predicted to have 3 TCR chains nor whole clones predicted to possess 3 TCR chains displayed an increase in doublet score (as defined by scDblFinder) compared to cells or clones with 2 TCR chains (**Extended Data Fig. 2c-d**). Higher doublet scores indicate a greater probability of being a doublet, so this suggests these clones are not enriched for doublets. Clones annotated to have 3 TCR chains also did not display elevated levels of RNA or unique genes (**Extended Data Fig. 2e-f**), again arguing against the notion that these 3 TCR chain clones are enriched for doublets. Rather, it is likely that technical dropout is the true cause of observing cells with fewer than 3 TCR chains within a VDJdive/ECLIPSE-defined clonotype that is deemed to truly express 3 chains. To test this notion, we performed TRUST4 reconstruction of TCR chains from the paired scRNA-seq gene expression data and identified TCR chains in both 2 and 3 chain clones that were predicted by ECLIPSE but not present in the TCR-sequencing library (**Extended Data Fig. 2g**). Overall, these analyses demonstrated that technical TCR chain dropout is occurring, and that VDJdive/ECLIPSE can mitigate its impact by recovering a portion of these lost chains. Our method thus results in higher fidelity clonal predictions that would not be possible by only considering the chains present in each cell.

### Validation of ECLIPSE in independent datasets

We next investigated whether VDJdive/ECLIPSE would improve clonotype predictions in additional, independent datasets across distinct biological contexts. For this analysis, we compared VDJdive/ECLIPSE against 3 widely used methodologies that were used as benchmarks in **Fig. 3e-g**, with each approach defining a clonotype by both CDR3 amino acid sequences but differing in the cells and chains that are filtered out. First, we compared ECLIPSE against these 3 approaches in a dataset of CD4+ and CD8+ T cells in patients with melanoma (Oliveira et al., 2021; Oliveira et al., 2022). In CD8+ T cells, ECLIPSE again displayed favorable performance when compared to these existing methods, decreasing TCR diversity (Shannon index) when compared to methods that consider all chains/cells as is and increasing TCR diversity when compared to stricter methods that remove cells/chains if the clonotype is ambiguous (**Fig. 5a**). Further, ECLIPSE substantially enlarged the average clone size for the 30 largest clones, with an over 100% increase in clone size in one patient when compared to the most restrictive standard method (**Fig. 5b**). We additionally confirmed that clonotype calling with ECLIPSE resulted in a substantial decrease in the number of T cells without an assigned TCR clonotype (**Fig. 5c**), supporting the ability of VDJdive/ECLIPSE to consistently enhance clone sizes and reduce missing data. We observed similar findings among CD4+ T cells from the same cohort of patients, although the increase in clone size was not as substantial (**Fig. 5d-f**). VDJdive/ECLIPSE therefore is highly useful for analyzing clonal dynamics within both CD8+ and CD4+ T cells.

**Figure 5:**
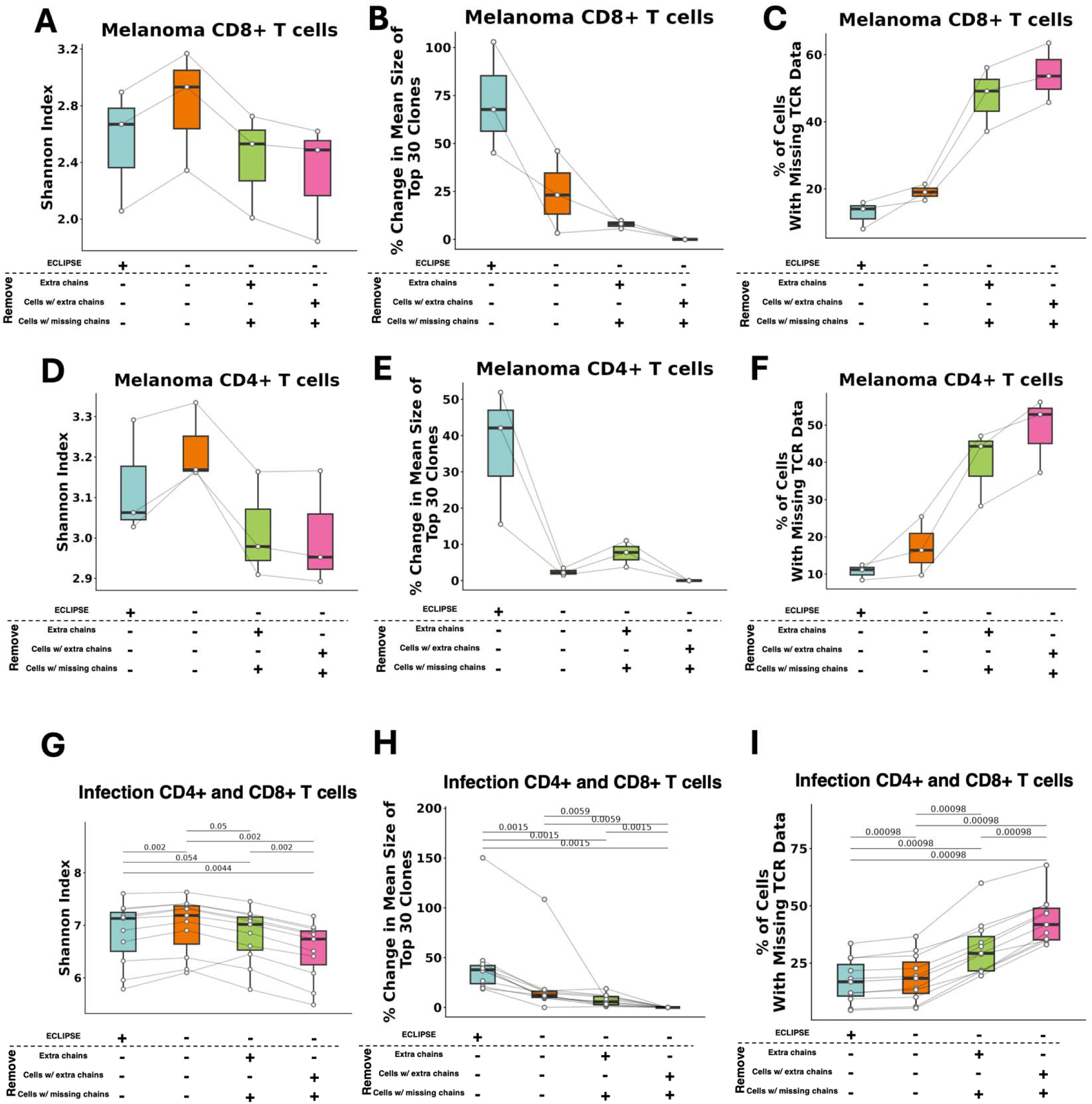
VDJdive/ECLIPSE are applicable to CD4+ and CDS+ T cells in diverse biological contexts. 3 validation cohorts. Row 1 CDS+ T cells in melanoma, row 2 is CD4+ T cells in melanoma (matched samples), and row 3 is patients with severe COVID-19 or severe bacterial pneumonia. Left (A, D, G) shows the Shannon index for TCR repertoire diversity per patient split by TCR analysis type. Middle (B, E, H) shows the percentage change in mean clone size of the 30 largest clones per patient split by TCR analysis type. Right (C, F, I) shows the percentage of cells without a clonal annotation per patient split by TCR analysis type. Wilcoxon-signed rank test was used for all plots with FDR correction being used for multiple-testing correction. p-values are shown above all applicable plots.

Last, we investigated the performance of VDJdive/ECLIPSE in a distinct biological context – infection. We analyzed a dataset of CD4+ and CD8+ T cells from the paired blood and bronchoalveolar lavage fluid (BALF) of 11 patients with severe COVID-19 or severe bacterial pneumonia (Zhao et al., 2021). In this context, Shannon repertoire diversity displayed a similar pattern as it did in both the settings of renal cell carcinoma and melanoma (**Fig. 5g**). As anticipated, this was accompanied by both an increase in clone size and a reduction in cells without a clonotype annotation (**Fig. 5h-i**). Ultimately, these findings indicate that regardless of disease context or T cell subset, VDJdive/ECLPSE substantially improve T cell clonal tracking, augmenting clone size and minimizing missing data.

## Discussion

scRNA/TCR-seq has revolutionized the tracking of individual T cell clones by dramatically increasing cellular throughput and speed. However, existing methods for translating recovered TCR chains to “true” clonotypes for each cell are limited. Clonotypes have classically been defined by the 1:1 pairing of CDR3 regions of TCR α:β, yet biological and technical factors drive many cells to possess more than 2 or only 1 TCR chain, rendering them clonally “ambiguous” (**Fig. 1b-c**) (Borcherding et al., 2020; Sturm et al., 2020; Zhu et al., 2023; Macosko et al., 2015; Qiu, 2020; Young and Behjati, 2020; Zheng et al., 2017; Malissen et al., 1988; Matis et al., 1988; Triebel et al., 1988; Padovan et al., 1993; Ji et al., *Nat. Immuno*., 2010). As a result, many pipelines filter out extra TCR chains based on expression level or entirely remove cells without a 1:1 α:β pairing, substantially reducing the already limited amounts of cells available for analysis, diminishing clone sizes, and distorting TCR diversity estimates (Sturm et al., 2020; Le Coz et al., *Sci. Immuno.*, 2023; Zheng et al., *Science*, 2021; Perez et al., *Science*, 2022; Carter et al., 2019; Muhowski and Rogers, 2023) (**Fig. 3e-g**, **Fig. 5a-i**). For these reasons, we propose that a novel methodology for TCR analysis is necessary, one that directly addresses both the biological and technical intricacies of scTCR-seq. Such a method ideally would maximize the accuracy and scale of clonal tracking while enabling users to make best use of the data generated from a relatively costly assay.

Here, we describe VDJdive, a computational method developed with these principles in mind; it implements the EM algorithm to predict the true clonotypes of ambiguous cells by considering known chain pairings from other cells in the same sample. We also introduced ECLIPSE (Enhanced CLonotypic Inference via Prediction of Single-cell Expression), a method which builds on VDJdive to discern true biological expression of extra TCR chains from technical artifacts and incorporate the presence of these 3 TCR chain clones into the final clonal annotations. Together, our approach greatly enhances the ability of users to track individual T cell clones through its ability to tangibly reduce the percentage of cells without annotated clonotypes and markedly increase the size of both moderate and large clones (**Fig. 3e-f, Extended Data Fig. 1a, Fig. 5b-c**, **Fig. 5e-f**, **Fig. 5h-i**). However, this increased power to annotate clones does not sacrifice accuracy: permutation testing revealed that, regardless of the chains detected in each cell, clones defined by VDJdive/ECLIPSE are far more phenotypically consistent than could be explained by chance (**Fig. 4e-f**). In fact, our approach is able to reliably predict TCR chains lost to chain “dropout” in the TCR-sequencing library but present in the cell and the RNA-sequencing library (**Fig. 4a, Extended Data Fig. 2g**). Similarly, it possesses the ability to decipher true biological expression of chains from false chains artificially added during data simulations (**Fig. 4b**). In this way, VDJdive/ECLIPSE hedge against the inevitable technical challenges of scTCR-seq (i.e. chain “dropout”, TCR contamination, cellular doublets), recovering the true biological signal present in each cell.

Our method also detected 3 TCR chain clones at a rate (patient median of 17.4% and 1.7% for TCRα and TCRβ, respectively) that was consistent with previous literature (**Fig. 3c**) (Muhowski and Rogers, 2023; Yang et al., 2020; Padovan et al., 1993, Carter et al., 2019). Patient variation was appreciable, with frequencies over 50% being observed in some patients. The percentage of cells with extra TCRα chains was also found to be positively correlated with the percentage of cells with extra TCRβ chains (**Fig. 3d**), perhaps suggesting that certain tumors may favor or select against T cells containing extra chains. Alternatively, some individuals may exhibit higher rates of allelic exclusion for both TCRα and TCRβ chains, impacting the average number of TCR chains per cell in the naïve T cell repertoire and before clonal expansion occurs. Previous literature has established that T cells with 3 TCR chains exhibit functional differences from classical 2-chain clones (Muhowski and Rogers, 2023). Specifically, it has been shown that CD4+ and CD8+ T cells expressing 3 TCR chains may exhibit increased expansion after antigenic exposure (either immunization or viral infection) than conventional cells, with CD4+ T cells but not CD8+ T cells remaining elevated in frequency at memory timepoints (Yang et al., 2020; He et al., 2002). This effect of heightened clonal expansion after antigen exposure was even found to be present in clones with 1 productive TCRα and 1 non-productive (out-of-frame) TCRα, suggesting that it is independent of potential dual antigen specificity that may be present through the expression of two chains (Dash et al., 2011). Given the ability of ECLIPSE to preserve 3 TCR clones with high fidelity, future studies will be able to use ECLIPSE to evaluate whether these described functional differences of 3 TCR chain T cells translate to humans and whether they impact disease progression or treatment efficacy. Such analysis cannot be performed with conventional methodologies that filter out cells with more than 2 TCR chains, making ECLIPSE a value tool for uncovering the biology of T cells with 3 TCR chains at a single-cell level.

More broadly, the reduced missing data and increased clone sizes provided by VDJdive/ECLIPSE will enhance clonal tracking for clonotypes of any chain count (2 or 3), particularly in humans. While studies utilizing adoptive transfer models in mice have been instrumental in developing our understanding of T cell differentiation and function, our understanding of such fundamental T cell processes is substantially more limited in humans (Gearty et al., 2022; Fagerberg et al., 2025). Paired scRNA/TCR-seq has emerged as a valuable alternative for human studies, with the RNA-sequencing library providing in depth phenotypic information for cells at each point in differentiation space and the TCR-sequencing library providing information on which cells share common ancestry. In this way, human T cell differentiation trajectories can be reconstructed, and this approach has proved fruitful in uncovering the distinct differentiation trajectories of tumor-reactive and bystander (non-tumor reactive) cells within tumors (Li et al., 2019; Zhang et al., 2018). The increased clone sizes generated by VDJdive/ECLIPSE will provide better statistical power for future research in this space, allotting improved ability to detect differences of between populations in a fixed total sample size. Likewise, the ability of VDJdive/ECLIPSE to mitigate the impact of technical challenges of scTCR-seq will reduce the noise associated with clonal tracking, leading to a similar effect. This increased accuracy may also support future research into the biological consequences of TCR repertoire diversity. Previous studies have reported improved overall survival in cancer patients with higher repertoire diversity. By providing more accurate diversity estimates, VDJdive/ECLIPSE may reveal additional consequences of TCR diversity (Hopkins et al., 2018; Keane et al., 2017).

In summary, VDJdive and ECLIPSE allow users to make the most informed use of their scTCR-seq data, resolving clonal ambiguity and hedging against technical artifacts of scTCR-seq. Their usage is broad and remains optimal in a diverse array of biological situations and T cell subsets (**Fig. 5a-i**). Furthermore, our approach is easily implementable: ECLIPSE operates with current standard workflows, namely Seurat for scRNA-seq analysis and scRepertoire for clonal visualization. Future studies with utilize VDJdive and ECLIPSE to uncover novel human T cell biology and further illuminate the role of T cells as both drivers of autoimmunity and mediators of protective immunity.

### Limitations

While VDJdive/ECLIPSE perform favorably against conventional TCR analysis methods in many contexts, the magnitude of benefit may not be always identical. Because VDJdive relies on known chain pairings in the sample to predict the clones of ambiguous cells, if the majority of cells in a sample are singletons, it will be unable to predict most ambiguous cells with high confidence. For instance, this could be the case with naïve or central memory cells in uninflamed lymph nodes or blood.

VDJDive (and as a result ECLIPSE) is also built on CDR3 amino acid sequences as the defining feature of a chain, but in rare circumstances two clones within a sample may exhibit identical CDR3 amino acid sequences with either a different nucleotide CDR3 sequence or a different set of variable, diversity, and/or joining gene segments. Users should also be cognizant that ECLIPSE specifically utilizes high confidence contigs from Cell Ranger’s all_contig_annotations.csv output, which differs from the filtered_contig_annotations.csv file that some other pipelines start with. In our opinion, high confidence contigs from the all_contig_annotations.csv file are ideal for clonal tracking as the increased number of chains elevates the precision of clonal tracking, however some clones that are annotated as a 3 TCR chain clone may include one chain that is either predicted by Cell Ranger vdj to be either non-productive, non-full length, or not from a real cell. Although not recommended for the purposes of clonal tracking, users can specify to ECLIPSE that they are only interested in contigs from the filtered_contig_annotations.csv file if they are cloning TCR chains and/or looking for the chains that are necessary for detecting a specified antigen.

## Methods

### VDJdive Methods

#### Basics and Code Availability

VDJdive is available as a package on <u>Bioconductor</u> and on <u>GitHub</u>. Also included on Bioconductor is a vignette demonstrating the use of the EM algorithm through the clonoStats function as well as other functionalities of VDJdive that were not discussed in this manuscript. These include diversity analysis and visualization.

#### Probabilistic Clonotype Assignment

Probabilistic clonotype assignment is performed in VDJdive with the clonoStats function. VDJdive takes as an input the set of filtered contigs assembled by the Cell Ranger vdj pipeline: filtered_contig_annotations.csv files. Clonotypes are defined as containing 1 TCRα chain and 1 TCRβ chain that are each identified by their CDR3 amino acid sequence. Cell barcodes with only a single, coherent pair of chains (i.e. 1 TCRα and 1 TCRβ) can be unambiguously assigned a clonotype label. In contrast, all other cells are considered to be clonally ambiguous.

In clonoStats, we implemented an expectation-maximization (EM) algorithm to probabilistically assign clonotypes to ambiguous cells (Dempster, Laird, and Rubin, 1977). The EM algorithm is widely used for imputing or assigning labels under partial uncertainty. Unlike other methods, for uncertain labels, they are assigned in a proportionally weighted manner that matches the confidence of the model that the label is correct (Do and Batzoglou, 2008). Similar methodologies have been successfully applied to estimate gene frequencies and to align multi-mapped sequencing reads (Ceppellini et al., 1955; Li and Dewey, 2011). In this context, we treat a cell’s true clonotype, represented by two specific chains, as a missing variable that can be inferred. For cells with ambiguous combinations of chains, we employ an EM algorithm to probabilistically assign clonotypes by leveraging information from other cells within the same sample. The objective is to maximize the likelihood function, *L*(θ), where θ represents the parameters of the marginal clonotype distribution. The observed data *x* includes the observed clonotype distribution for unambiguous (i.e. clonally resolved) cells and for each ambiguous cell, the set of clonotypes with which the observed chains would be consistent. The missing information is modeled as a latent variable *z*, representing the true, but unobserved chain pairings/clonotype for each cell.

In the E-step, we compute the conditional expectation of the complete-data log-likelihood, given the observed data *X* and the current estimate of the parameters θ^(t)^, as follows:

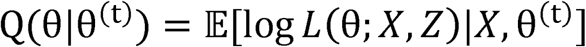

In practice, this represents the proportional assignment of labels to ambiguous cells, based on the relative abundance of each possible clonotype. For initialization, ambiguous cells’ labels are split evenly amongst all possible clonotypes with which their observed set of chains is consistent. In the M-step, we maximize this expected complete-data log-likelihood to update the parameters *θ*:

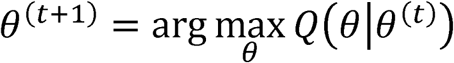

This step refines the estimates of the marginal clonotype distribution, based on the partial assignments calculated in the E-step. These steps are repeated iteratively until the parameters converge, meaning that the maximum change in any element of *θ*^(t)^ between successive iterations is less than a pre-defined threshold, i.e. 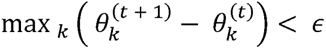, where *k* indexes all possible clonotypes in the data.

After convergence, VDJdive utilizes the final estimated clonotype distribution parameters (*θ*) to calculate the final probability of each clonotype being the true clonotype for each cell. The probabilities also represent the proportion of partial assignment, meaning that as the probability of each clonotype being true increases, so too does the proportion of the cell that is assigned to the clonotype. The data can then be returned in an assignment matrix, where the probability of each clonotype being true for each cell is documented.

### ECLIPSE Methods

#### Code Availability

ECLIPSE is an R package that will be available as a package at the time of publication and on <u>GitHub</u>. While most functionalities of ECLIPSE were discussed here (implemented through tcrEclipseMulti), it also includes a function (tcrDoubletDetect) that should be run after running tcrEclipseMulti which allows the user to remove cells with >4 total (or >2 of a type) TCR chains detected and set an appropriate threshold for the number of unique UMIs based on cells that express high numbers of TCR chains.

#### Contig Importing and Filtering

By default, ECLIPSE uses all_contig_annotations.csv files from the output of Cell Ranger vdj, although users can specify if they would like to use filtered_contig_annotations.csv files instead. From there, only high confidence contigs with cell barcodes that are found in the provided Seurat object are kept. Contigs must have a CDR3 sequence present that is ≤ 30 amino acids in length without a stop codon inside the CDR3 region (as indicated by a * by Cell Ranger vdj).

Any contig that has a TRDV segment joined to a TRAJ segment as well as a TRAC segment is reannotated to be a TCRα chain. This is because some TRDV segments, particularly TRDV1 in humans, can join with TRAJ/TRAC segments to form productive TCRα chains that pair with TCRβ chains (Volkmar et al., 2023; Miossec et al., 1990; Heather et al., 2025). However, Cell Ranger labels such chains as either TCRδ or of multiple types, making manual adjustment to TCRα necessary. After this adjustment, all contigs that are not TCRα or TCRβ are removed. Lastly, any duplicated contigs (defined here as contigs that have the same cell barcode, chain type, and CDR3 amino acid sequence) are filtered to retain only 1 copy per cell.

#### Clone Calling

Filtered contigs are fed into a custom function described in a later section: findSpecialClones. This function detects and preserves clones with 3 TCR chains from being coerced into having 2 by the EM algorithm. findSpecialClones is run twice, with the first instance working on clones with 2 TCRα and 1 TCRβ, and the second working on clones with 1 TCRα and 2 TCRβ. After these clones are modified, they are provided to the clonoStats function of VDJdive where the EM algorithm is implemented (using the arguments method = “EM”, type = “TCR”, assignment = TRUE, and group). This generates a matrix providing the probability of each clonotype for each cell (equivalent to the proportion of each cell assigned to each clonotype), from which the clone with the highest probability (*C_1_*), its probability (*p*_1_), and the probability of the 2^nd^ most likely clone (*p*_2_) are extracted. The ratio (*r*) of the highest probability over the 2^nd^ highest probability is also calculated, where 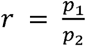. Cells are fully assigned by ECLIPSE to one clone if one of the following thresholds is met:

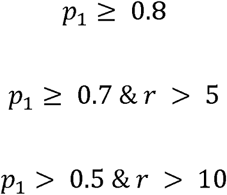

In this way, cells that are very likely (e.g. *p*_1_ = 0.995) to be in a clone are assigned. Cells that are less likely, but with one clear best option (*r* > 5 - 10) can also be assigned given that ratios that are that high generally indicate clonotypes other than *C*_1_ are noise. For example, there can be cells where *p*_1_ = 0.75 and *p*_2-*n*_ are all < 0.001. As an additional filtering step, all cells that meet one of these thresholds must not also have contained only a singular contig that was only seen in that cell. If a contig was seen alone and only in one cell, the cell is treated as if none of the thresholds were met.

For all cells where the thresholds are not met, no prediction is made. Instead, the contigs are considered as is based on CDR3 amino acid sequences of the present chains, in that the final clonotype can contain only 1 chain, 2 chains, or more than 2 chains. Additionally, for clones that have 3 total chains, any cell containing or predicted by VDJdive to contain 2 of the 3 chains (with 1 bitypic and 1 monotypic chain) is treated as having all 3 chains.

#### Detecting Clones with Extra TCR Chains

The findSpecialClones function is used by ECLIPSE to detect and preserve clones that likely contain 3 TCR chains. This is important because the EM algorithm of VDJdive assumes all cells must possess 1 TCRα and 1 TCRβ, so separate retention of clones with 3 TCR chains is necessary. The main requirement by findSpecialClones for calling a clone as having 3 TCR chains is 3+ cells having the same 3 TCR chains. Alternatively, 1+ cell with all 3 chains, 1+ cell with one of the bitypic chains alongside the monotypic chain, and 1+ cell with the other bitypic chain alongside the monotypic chain. If 1 of these 2 thresholds is met, it is very likely that the clone is biologically true, given that doublets or TCR contamination would be unlikely to repeat in the same way in multiple cells.

There are also additional filtering steps to ensure that no doublets or incidents of TCR contamination result in a clone being called with 3 TCR chains. The first is that for every monotypic chain of a clone with 3 TCR chains, ≥ 10% of all cells in the sample with the monotypic chain must possess both paired bitypic chains. Additionally, both bitypic chain types must be most commonly paired in that sample to the monotypic chain.

After the selection of clones with 3 TCR chains, cells missing the monotypic chain but potentially in each of these clones are considered. For each potential clone, if ≥ 80% of cells across the sample with some combination of the bitypic chains contains the same set of monotypic and bitypic chains, then other cells potentially of the same clone with 1-2 bitypic chains and no monotypic chain are assigned to have the bitypic chain set of this dominant subset. This is necessary because it allows the EM algorithm to consider 3 TCR chain clones for these cells that are missing a monotypic chain.

#### Integration of TCR Data with Seurat Object and Downstream Visualization

For each cell, the final clonotype (either predicted or just the chains present) is added to the Seurat object that was originally provided by the user (Hao et al., 2023). Additionally, regardless of whether a prediction is made, other columns are added by the combineTCR and combineExpression (with cloneCall = “aa” and proportion = TRUE) functions of scRepertoire to provide information about the original contigs that were present (Borcherding et al., 2020; Yang et al., 2025). This includes the gene segments, CDR3 amino acid sequences, CDR3 nucleotide sequences. In this way, the user can compare the predicted clonotype against the original chains that were detected for each cell.

For each clone, the proportion, clone size, and relative clone size (i.e. small or hyperexpanded) are also calculated in the Seurat object, and this is done in a way that formats the Seurat object for downstream use with most functionalities provided through scRepertoire. Examples of this include diversity calculations (clonalDiversity), tracking clones across tissues/groups (e.g. clonalCompare), and visualization (e.g. clonalHomeostasis). Importantly, because ECLIPSE utilizes amino acid CDR3 sequences, scRepertoire analysis must include the argument cloneCall = “aa”. Also present in the final Seurat object provided to the user is a column that notes if the cell is predicted to be a MAIT cell or iNKT cell based on TCR segment usage.

### General Methods

#### General Analysis and Visualization

All analysis was performed in R version 4.4.1 (R Core Team, 2024) using RStudio (Posit team, 2025). Data wrangling was performed with the tidyverse package (Wickham et al., 2019). ggplot2 was used for producing graphs, with ggsci and RColorBrewer being used for color palettes. (Wickham, 2016; Xiao et al., 2025; Neuwirth et al., 2022). BioRender was used to create the integrated workflow graphic.

#### Artificial Intelligence Usage Statement

Small amounts of R code were drafted with the use of ChatGPT. However, all code was manually reviewed and edited by the authors, who take full responsibility for all code present. All text was human written.

#### Statistics

All significance testing was performed in R. ggpubr (Kassambara, 2025) was used for comparing 2 groups or correlation tests, specificially with stat_compare_means or stat_cor. For tests of more than 2 groups, repeated Wilcoxon signed rank tested were performed with wilcox.test. To correct for multiple-testing, p-values were altered using p.adjust for FDR correction. ggpubr was then used to add the p-values to each graph.

#### Data and Code Availability

All code used in the generation of plots/analysis will be available at the time of publication on GitHub.

Most analysis (except **Fig. 5**) was performed on a scRNA/scTCR-seq dataset of paired samples of tumor and adjacent normal tissue from 13 patients with clear cell renal cell carcinoma (ccRCC) (Braun et al., 2021). This data is available on dbGaP with the accession number phs002252.v1.p1. Alternatively, a post-quality control filtering counts matrix is available as supplemental data in the original manuscript, as well as cell metadata. Importantly, for all analysis in this manuscript, the TCR FASTQ files were realigned with the current version of Cell Ranger vdj at the time of analysis (9.0.1) using the Human V(D)J 10X Genomics GRCh38 Ensembl 7.1.0 reference.

For the melanoma datasets, all patients had stage III/IV disease, with the dataset containing 5 tumor samples from 4 patients (Ott et al., 2017; Oliveira et al., 2021; Oliveira et al., 2022). However, only 3 patients were considered for our analysis since one patient (Pt-B) had low cell count. The raw data is available on dbGaP with ID 26121 and accession phs001451.v5.p1. The Seurat objects used for analysis were generated as previously described in the original manuscripts (Oliveira et al., 2021; Oliveira et al., 2022).

For the COVID-19/bacterial pneumonia dataset, data is available on Gene Expression Omnibus under the accession number GSE167118 (Zhao et al., 2021). Here we used CD4+ and CD8+ T cells, and the samples are from the paired blood and bronchoalveolar lavage fluid (BALF) of 11 patients with severe COVID-19 or bacterial pneumonia.

#### TCR Repertoire Diversity Analysis

Repertoire diversity with calculated using the clonalDiversity function of scRepertoire (Borcherding et al., 2020; Yang et al., 2025).

#### TCR Chain Removal or Addition Simulations

For chain removal simulations, we began by running the standard ECLIPSE filtering as described above. Among cells with 1 TCRα and 1 TCRβ, we randomly selected 500 contigs (each from a different cell) and removed them from the contigs. As 55,957 cells had at least 1 TCR chain, this represented nearly 1% of all cells. All other cells in the dataset were unmodified. Only cells with 1 TCRα and 1 TCRβ were modified as this represented a ground-truth clonotype, whereas cells other chain configurations were clonally ambiguous.

After contig removal, the 2^nd^ half of the standard ECLIPSE pipeline was run as previous described. The clonotypes from pre-removal were compared to the clonotypes from post-removal for the 500 altered cells to see if VDJdive/ECLIPSE can mitigate the impact of chain dropout and do so accurately. 3 possibilities were considered: the call was the same, there was a new (likely incorrect) clonotype, or no prediction was made due to the contig removal (indicating VDJdive/EM algorithm did not assign any 1 clone with high confidence and instead defined the cell by just 1 chain). As a control, we also performed an identical comparison of whether the clonotype changed for the ∼99% of cells that were unmodified in the simulation. As these cells were not modified (only other cells in the same sample were), the final clonotype should not be impacted.

For chain addition simulations, it began similarly, however, among cells with 1 TCRα and 1 TCRβ, 500 barcodes were randomly selected for chain addition, and 500 contigs were selected from 500 other cells. The selected contigs were duplicated, and each was assigned to one of the 500 target barcodes. For this, their barcode, sample, sequencing run, and contig_id were altered to that which would match the new barcode. The result was 500 random additional contigs added to 500 random cells with a ground-truth clonotype. After this addition, an identical protocol to the chain removal simulation was followed.

#### Permutation Tests

For the permutation test of **Fig. 4e**, the “major” and “minor” bitypic chains of each 3 TCR clone were determined by analyzing how many cells of the clone each chain was detected in. Additionally, for each set of chains detected for each clone, the percentage of cells in each of the 13 clusters was calculated.

Using this percentage information, the total variation distance of cluster usage for cells expressing the bitypic major vs. the bitypic minor chain was found. The calculation of total variation distance is given by the following equation:

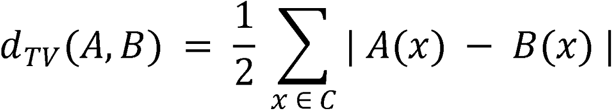

where *d_TV_* (*A*, *B*) is the total variation distance of cluster usage between cell populations *A* and *B, C* is all existing clusters (13 for the renal cell carcinoma dataset), and *A(x)/ B(x)* represent two probability mass functions equal to the probability of cells in either cell population A or B falling into each cluster (here bitypic major expressing cells vs. bitypic minor expressing cells). 0 ≤ *d_TV_*(*A*, *B*) ≤ 1, where *d_TV_*(*A*, *B*) = 0 represents perfectly overlapping cluster usage, and *d_TV_* (*A*, *B*) = 1 represents perfectly distinct cluster usage.

After calculating the total variation distance for each clone between cells that express the bitypic major vs. minor chain, the mean total variation distance across all 3 TCR clones was calculated. We then computed the total variation distance for random sets of clones with 1 TCRα and 1 TCRβ compared to other random clones. Each comparison was between clones of the same donor. The number of clones in each set was equal to the number of 3 TCR clones among all donors, and the mean of each total variation distance was found, giving a randomly permuted mean total variation distance.

After repeating this calculation of a randomly permuted mean total variation distance for a total of 10,000 times, we calculated a permuted p-value by finding the number of times the observed mean value was less than or equal to a permutated mean value and dividing that number by 10,000 (the number of permutations). Under a null hypothesis where VDJdive/ECLIPSE combine biologically distinct cells to form clones, the observed mean total variation distance should fall in the distribution of permuted mean total variation distances.

For the permutation test of **Fig. 4f**, the only difference was the compared groups were cells missing either TCRα or TCRβ vs. those that contained both chains and were called in the same clone. The permutation test assessed random sets of cells with only TCRα or TCRβ against random sets of cells with both chains.

#### Doublet Score Calculation

All cells sequenced from the original study (not just CD8+ T cells) were run using scDblFinder using the following arguments: samples, BPPARAM = MulticoreParam(3), clusters, and dbr = 0.001) (Germain et al., 2022). A relatively low doublet rate was chosen as doublet removal had already been performed on the object during the original study, so few doublets were expected to be remaining. Using the output, we calculated the median doublet score in CD8+ T cells per group and compared the groups.

#### TRUST4 TCR Reconstruction

TCR reconstruction from the RNA library for each sample was ran using TRUST4, allowing for validation of what was predicted to be missing by VDJdive/ECLIPSE by comparing against some of what was confirmed to be missing from the TCR library (Song et al., 2021). Arguments used include t (8), 1 and 2 (specifying the files), f (path to fasta file), ref (providing the TCR reference sequence, the Human V(D)J 10X Genomics GRCh38 Ensembl 7.1.0 reference), barcode (path to the R1 file), readFormat (bc:0:15), o (output file prefix), and od (directory for output files).

After TRUST4 was run, the airr output files were imported into R. Each reconstructed chain was compared to what was predicted by VDJdive/ECLIPSE and what was detected by sequencing through the TCR library.

#### Research Ethics Statement

No new human samples were generated in this study. All datasets used here were previously published and received de-identified, with all previous studies having IRB approval. As such, all research in this study has followed ethical standards relating to human research. No animals or datasets from animals were used in this study.

## Acknowledgments

We would like to thank members of the Braun and Street labs for fruitful discussions that have helped to improve ECLIPSE. We are particularly grateful for Dr. Zachary Yochum, Colin Laughlin, and Nelson Moritz of the Braun lab. We also would like to thank the Irizarry lab (in particular Linglin Huang and Jared Brown) for their conversations on single-cell sequencing and EM algorithm optimization. Additionally, we are grateful to Dr. Gwo-Shu Mary Lee for her advice. Lastly, we would like to thank Lena Wirth for allowing us to test ECLIPSE on her dataset and reviewing the manuscript.

## Funding

E.C.B. is funded by a T32 (T32AI007019-50S1) from NIAID/NIH and would like to thank the Gruber Foundation for support (Gruber Science Fellowship). D.A.B. acknowledges support from the DOD (KC190128, KC220016, KC240196), the Cancer Research Institute (CRI), the NIH/NCI (1R37CA279822-01), the Louis Goodman and Alfred Gilman Yale Scholar fund, and the Yale Cancer Center (supported by NIH/NCI research grant P30CA016359). K.S. and M.Y. received funding from the NIH (P01CA196569).

## Disclosures

D.A.B. reports share options in Elephas; advisory board, consulting or personal fees from Cancer Expert Now, Adnovate Strategies, MDedge, CancerNetwork, Catenion, OncLive, Cello Health BioConsulting, PWW Consulting, Haymarket Medical Network, Aptitude Health, ASCO Post and Harborside, Targeted Oncology, Merck, Pfizer, MedScape, Accolade 2nd MD, DLA Piper, Dechert, AbbVie, Compugen, Link Cell Therapies, Scholar Rock, NeoMorph, Nimbus, Exelixis, AVEO, Eisai, Daiichi Sankyo, Caris, and Elephas; and research support from Exelixis and AstraZeneca, outside of the submitted work. All other authors report no relevant disclosures.

---

**Extended Data Figure 1:**
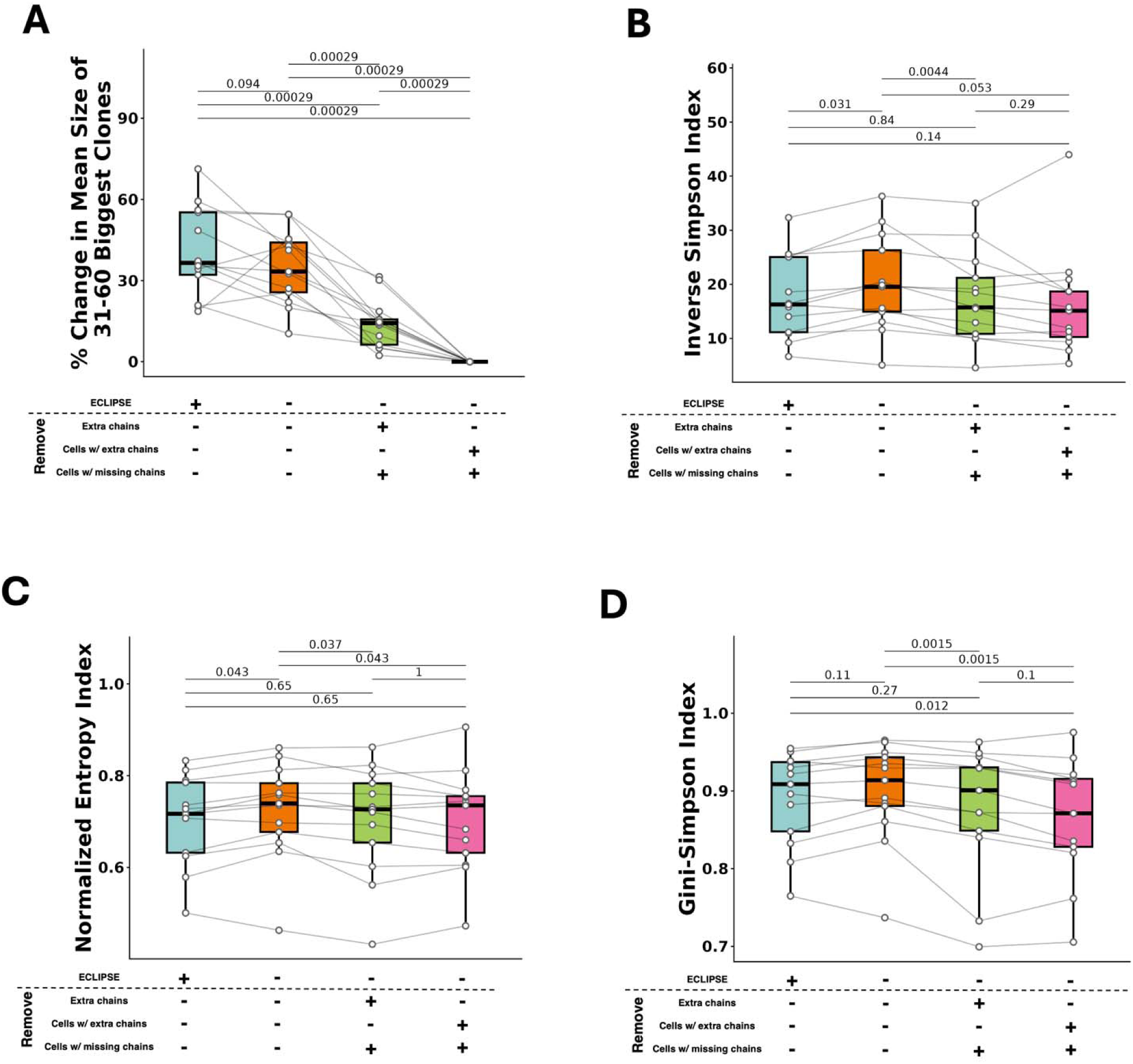
VDJdive/ECLIPSE enhance clone sizes and refine TCR repertoire diversity estimates. A. Percentage change in mean clone size for the 31st-60th largest clones (mean size of 8 cells) per patient in the renal cell carcinoma dataset. This is split by TCR analysis type. B. TCR diversity per patient as measured by the Inverse Simpson index and split by TCR analysis type. C. Identical to B, but with the Normalized Entropy index. D. Identical to B-C, but with the Gini-Simpson index. Wilcoxon-signed rank test was used for all plots with FDR correction being used for multiple-testing correction. p-values are shown above all applicable plots.

**Extended Data Figure 2:**
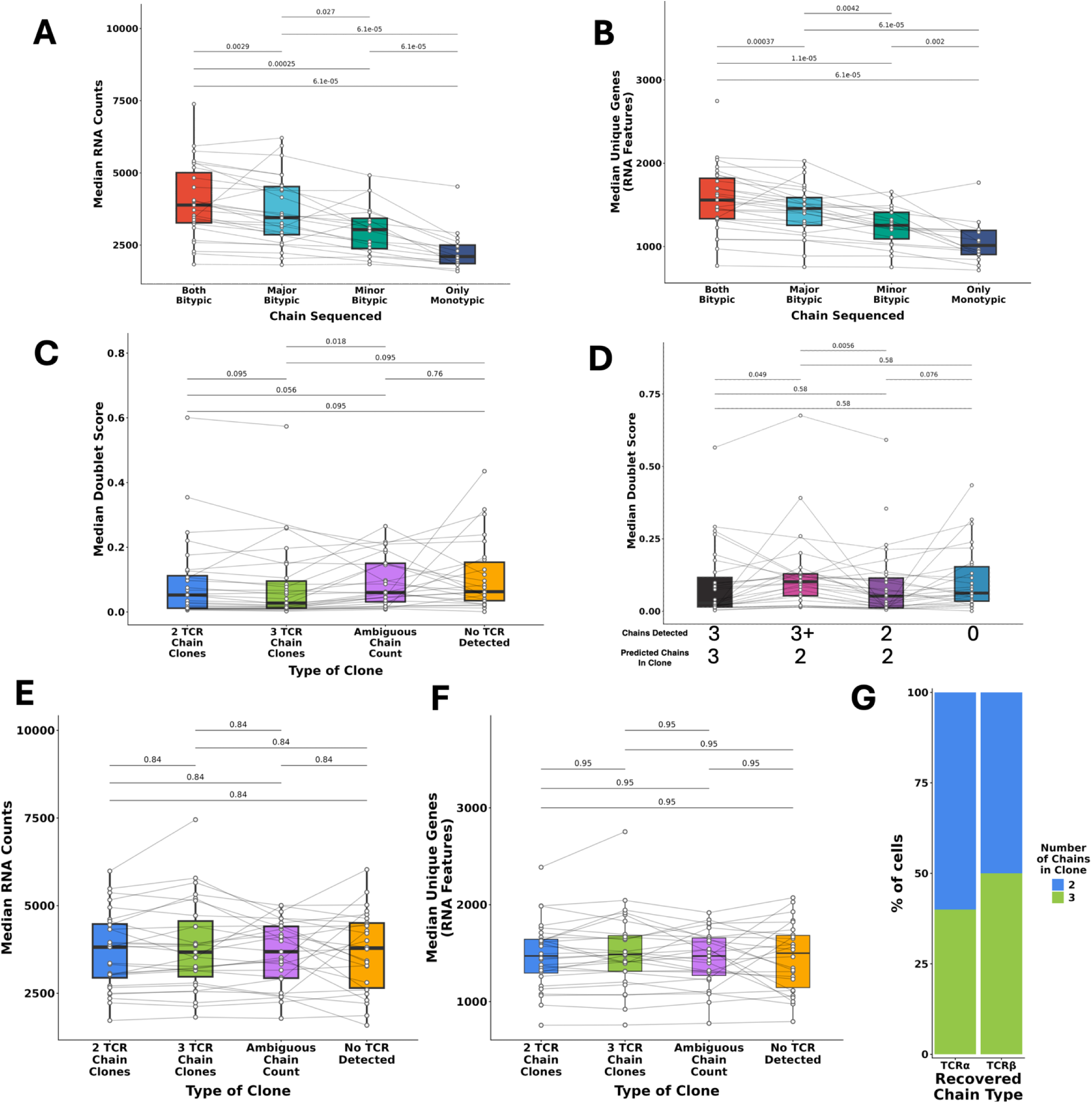
Clones with 3 TCR chains are not enriched for doublets and instead technical artifacts reduce the number of TCR chains detected in many cells. A. For 3 TCR chain clones, the median RNA counts per sample is shown split by the bitypic chains that were detected. B. Same as A, but with median unique genes. C. The median doublet score per sample is shown, split by the number of chains in the clone. D. Same as C, but split by the number of chains detected and the predicted number of chains. E. The median RNA counts per sample split by clone chain count is shown. F. Same as E, but with median unique genes. G. TRUST4 reconstruction of TCRs from the RNA-sequencing library. The graph shows TCRs that were predicted by VDJdive/ECLIPSE and not present in the TCR-sequencing library, but confirmed by TRUST4 to have been present in the cell (i.e. the RNA-sequencing library). The colors show the number of predicted chains in the clone of the recovered chain. Wilcoxon-signed rank test was used for plots A-F with FDR correction being used for multiple-testing correction. p-values are shown above all applicable plots.

